# Dynamic regulation of a domesticated DNA transposon-derived gene, in human germline and early embryonic development

**DOI:** 10.1101/2025.10.17.683089

**Authors:** Tanuja Bhardwaj, Sharmistha Majumdar

## Abstract

Embryonic development requires precise transcriptional regulation to guide transitions from totipotency to pluripotency and lineage specification. Transposable element (TE) derived genes are increasingly recognised as regulators of these processes, but the role of DNA transposon-derived factors remains poorly understood. Here, we investigate human embryogenesis, primordial germ cell (PGC) development, and gametogenesis, using bulk and single-cell transcriptomic and epigenomic datasets. We observe that THAP9, a domesticated DNA transposable element-derived gene, is selectively enriched in PGCs, but transiently silenced in somatic lineages. Moreover, the gene exhibits lineage-specific expression during pre-implantation development, peaking at the zygote stage, transiently paused during cleavage, reactivated at the morula, and remains enriched in trophectoderm and embryonic stem cells but repressed in primitive endoderm and epiblast. Interestingly, THAP9 is dynamically regulated across gametogenesis, with prominent expression in spermatids and germinal vesicle oocytes. Epigenomic profiling revealed progressive chromatin remodelling, including early bivalency and enhancer activation, consistent with roles in transcriptional reprogramming. These findings identify THAP9 as a germline-enriched, developmentally regulated gene with sex-specific functions in gametogenesis and pluripotency, advancing our understanding of how domesticated transposon-derived genes contribute to human reproduction and embryogenesis

## Introduction

Embryonic development is a tightly controlled process involving a network of genes that ensures that a single cell develops into a functional organism [1] via tightly regulated cell division, proliferation, and differentiation. As cells progress from totipotency to pluripotency and continue to their fate determination, including the zygotic genome activation (ZGA)[2] and development of primordial germ cells (PGCs), which itself retains its potential to form a new organism [3], a series of changes occur in the expression of various genes, including transposable elements (TE) [4]. TEs, or mobile elements, comprise ∼53% of the human genome [5]. Transposable elements play a significant role in the evolution of the human genome. Many TEs have been domesticated (lost mobility) to perform different functions [6]. TEs often regulate genes through their action as cis-regulatory elements supplying promoters, enhancers, and boundary elements, hence playing an important role in type-specific expressions within cells [7]. TEs express themselves in a highly regulated stage-specific manner [5] during mammalian embryogenesis and play crucial roles in transcriptional regulation [8], X-chromosome inactivation [9], and pluripotency maintenance [10]. This regulatory function is particularly noteworthy in pluripotent stem cells, where TEs can alter the transcription levels of genes required for their maintenance of pluripotency and also determine cell fates [11], and can significantly affect embryo development and male infertility by influencing sperm function, epigenetic modifications [12], and overall reproductive outcomes[13]. Understanding the role of transposable elements in these processes is essential for understanding embryonic development and addressing issues related to fertility.

THAP9 (Thanatos-associated protein 9) is a gene derived from a DNA transposable element that is “domesticated” in the human genome [14]. THAP9 is a member of the THAP protein family, characterised by the presence of a DNA-binding domain, called the THAP domain [15]. Members of this family function as transcription factors [16,17], with several involved in controlling the expression of genes regulating angiogenesis and apoptosis [18], cell cycle control [19] and epigenetic gene silencing. THAP proteins have been implicated in various human diseases, including cardiovascular disorders, DYT6 dystonia, and cancer [15,20,21]. Notably, THAP11 regulates the pluripotency of embryonic stem cells [22], highlighting the developmental significance of this protein family. Despite its membership in this unique family of proteins, the exact cellular function and physiological role of THAP9 are still unknown. Our studies in oligodendrocyte lineage cells indicate that THAP9 is upregulated during the transition from oligodendrocyte progenitor cells (OPCs) to myelinating oligodendrocytes and is co-expressed with members of the SOX family and other developmental regulating genes [23]. However, the absence of THAP9 homologues in standard model organisms such as mice and zebrafish has limited investigation of its potential function in human development, representing a significant gap in our understanding of human developmental biology.

The unique evolutionary origin of THAP9 as a “domesticated” DNA transposase-derived gene presents an exceptional opportunity to understand how transposable element-derived genes contribute to human-specific aspects of development and reproduction. While numerous studies have explored the role of retrotransposons in embryonic development and germ cell biology, much less is understood about the contribution of DNA transposons to these processes. Given that, we hypothesise that THAP9 may play a significant regulatory role in human embryogenesis and primordial germ cell (PGC) development.

This study aims to address this knowledge gap by comprehensively characterising the THAP9 gene role in human germ cell development and early embryogenesis. We sought to determine whether THAP9 functions in naive cellular states and during differentiation processes, similar to our observations in oligodendrocyte development. Specifically, we analysed the transcriptional landscape of THAP9 in human germ cells and pre-implantation embryos, examined its epigenetic regulation, and identified potential downstream targets and regulatory networks.

Spermatogenesis is a continuous process that occurs throughout adult life, producing four motile haploid spermatozoa from each diploid spermatogonial precursor, whereas oogenesis is a discontinuous and temporally extended process that results in a single large, non-motile oocyte, characterised by prolonged meiotic arrest from fetal development until ovulation. Given the fundamental differences between spermatogenesis (germ cell maturation in males) and oogenesis (germ cell maturation in females), particularly their distinct temporal patterns and regulatory requirements, investigating THAP9 expression and function during these developmental stages will provide essential insights into how “domesticated” transposon-derived genes coordinate multiple aspects of human reproductive biology and early development. These contrasting cellular outcomes and timelines highlight the distinct regulatory environments within which THAP9 may function and mechanisms that may govern its activity during gametogenesis.

## Results

THAP9 exhibits restricted expression patterns in adult human tissues, with notably low expression *(<5 TPM)* across most somatic tissues according to GTEx bulk RNA-seq data [24] (Suppl. Fig. 1). However, THAP9 shows detectable expression in the testes and notable expression in oocytes (Suppl. Fig. 1), indicating its potential involvement in both male and female reproductive biology. Interestingly, the second-highest expression is observed in the thyroid, an endocrine organ that plays a key regulatory role in reproductive function, further supporting a link between THAP9 activity and reproductive physiology. This tissue-restricted expression pattern, combined with our previous observations of THAP9 upregulation during oligodendrocyte differentiation [23], led us to hypothesise that THAP9 may function specifically during cellular differentiation processes, particularly in reproductive and developmental contexts. To investigate the potential role of the THAP9 gene in human development and reproduction, we comprehensively analysed its expression patterns across multiple developmental stages and cell types. We examined THAP9 expression during pre-implantation embryonic development, primordial germ cell (PGC) specification and development (weeks 4-19), adult spermatogenesis, and oogenesis. Additionally, we analysed associated epigenetic modifications to understand the regulatory mechanisms governing THAP9 expression in these developmental contexts.

### THAP9 exhibits a distinct cell-type-specific expression pattern during human embryonic development

To investigate the developmental expression patterns of THAP9, we first examined its expression during human embryonic development using single-cell RNA-seq data (Table) from primordial germ cells (PGCs) and somatic cells across developmental weeks 4-19 (Figure 1a). This analysis revealed that THAP9 exhibits predominant expression in PGCs compared to somatic cells across all developmental timepoints examined (Figure 1b). THAP9 expression was consistently higher in PGCs than somatic cells throughout development in both sexes, with particularly striking differences at weeks 7 and 8-11 (p<0.0001 and p<0.001, respectively), suggesting a specialised role in germline biology.

**Figure 1:**
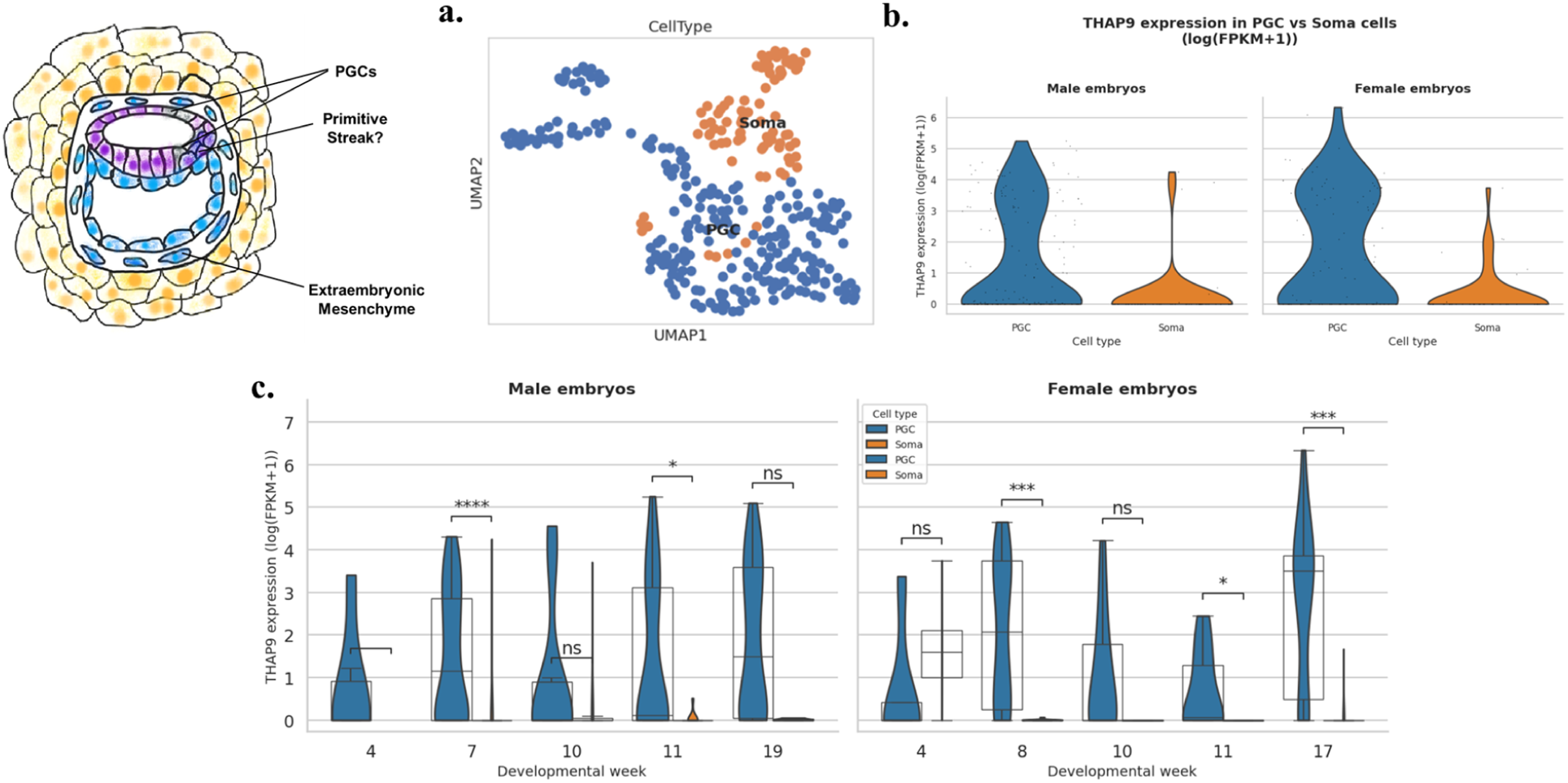
THAP9 expression analysis across embryonic development shows distinct cell-type specificity. **(a)** UMAP visualisation of single-cell RNA-seq data showing clear separation between primordial germ cells (PGCs, blue) and somatic cells (Soma, orange) based on transcriptomic profiles. **(b)** Overall, THAP9 expression distribution comparing PGCs (blue) versus somatic cells (orange) in male (left) and female (right) embryos **(c)** Developmental time-course analysis of THAP9 expression (log(FPKM+1)) in male (left) and female (right) embryos from weeks 4-19 and 4-17, respectively. Violin plots show expression distributions with overlaid box plots indicating median and quartiles. Statistical comparisons between PGCs and somatic cells at each time point were performed using the Mann-Whitney U test (*p<0.05, ***p<0.001, ****p<0.0001, ns = not significant).

Notably, somatic cells demonstrated progressive downregulation of THAP9 expression during development, with minimal to undetectable expression by later developmental stages (Figure 1c). This temporal decline indicates that THAP9 function may be restricted to early embryonic processes in somatic lineages. In contrast to somatic cells, PGCs sustained robust THAP9 expression throughout the examined developmental window, suggesting continued functional importance in germline maintenance, migration, or early gametogenesis. Expression levels in PGCs showed minimal sex-specific differences, indicating that the role of the THAP9 gene in germline functions is likely conserved between male and female germ cells during these early developmental stages. The highest PGC expression occurred around weeks 7-11 (Figure 1c), coinciding with critical periods of germline specification and early gonadal development, further supporting a role in fundamental germline processes.

These findings establish THAP9 as a germline-enriched gene during human embryonic development, setting the foundation for investigating its continued role in adult gametogenesis and early embryonic development.

### THAP9 is transcriptionally activated during human preimplantation development through coordinated chromatin remodelling

Given the strong germline expression observed during embryonic development, we next investigated whether THAP9 plays a role in the earliest stages of human development by analysing its expression dynamics during preimplantation development (Figure 2a). We examined transcriptomic data from 73 embryos representing six developmental stages (Table 1): zygote (1-cell, n = 13), 2-cell (n = 13), 4-cell (n = 16), 8-cell (n = 15), morula (n = 3), and blastocyst (n = 13). Examination of THAP9 expression revealed minimal transcript levels in the 1C, 2C, and 4C stages, followed by a significant increase at the 8-cell to morula transition (log2FC = 0.84 and 1.7, respectively; p < 0.05) (Figure 2b). Expression levels then plateaued in the blastocyst stage (log2FC = 0.95, not significant) (Figure 2b). These transcriptional changes coincided with stage-specific epigenomic remodelling at the THAP9 locus, providing mechanistic insight into its developmental regulation. By this developmental stage, the embryo has undergone lineage segregation into the trophoectoderm (TE) and inner cell mass (ICM) (Figure 2a), from which the primitive endoderm and epiblast subsequently emerge.

**Figure 2:**
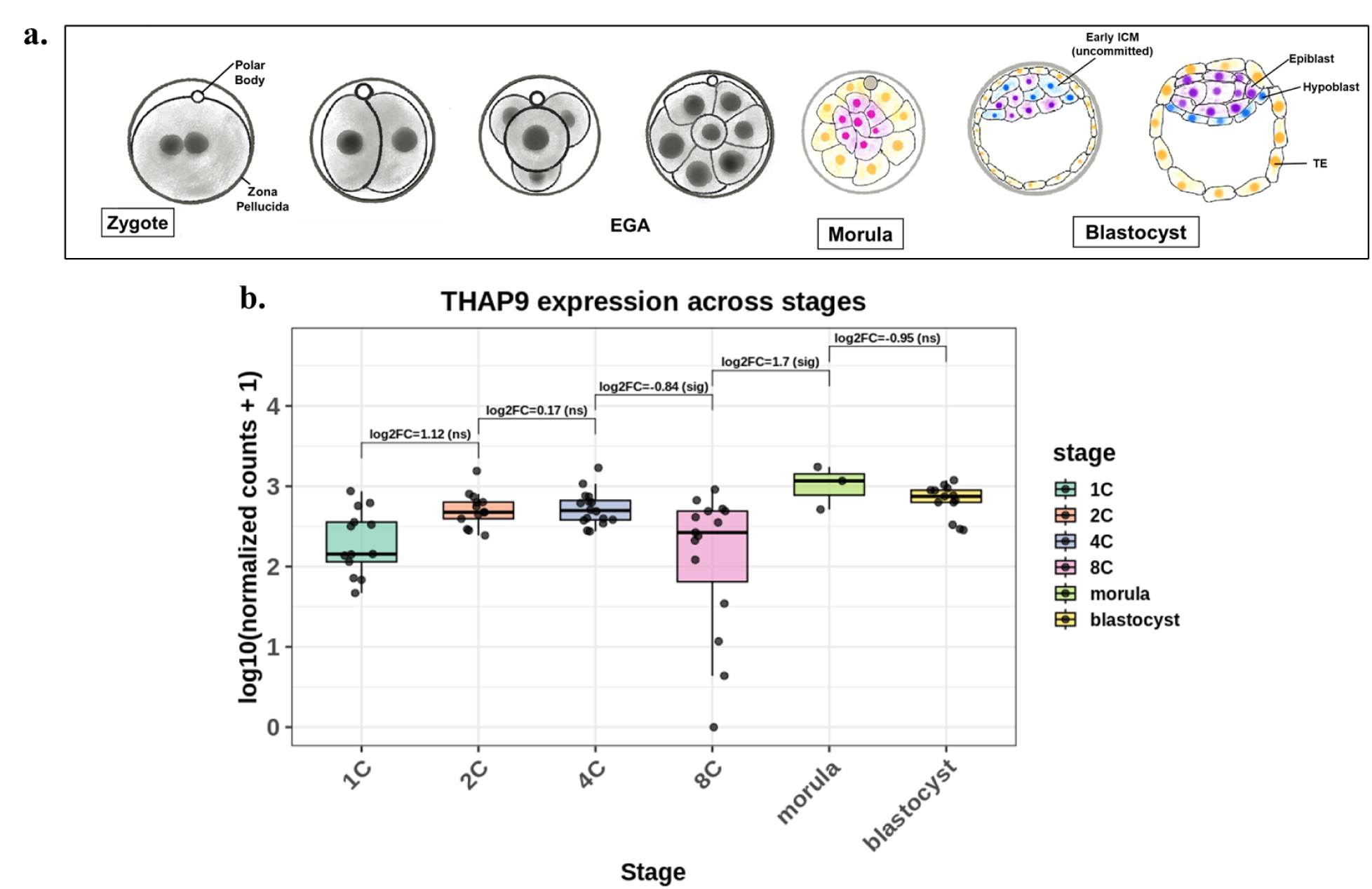
Stage-specific expression and regulation of THAP9 during human preimplantation development: **(a)** Schematic of developmental stages analysed from zygote to blastocyst (1C, 2C, 4C, 8C, morula, blastocyst), highlighting key cell lineages. **(b).** THAP9 expression pattern across embryonic developmental stages. Boxplot displaying THAP9 (ENSG00000168152) expression levels as log10-normalised counts+1 across six consecutive stages of pre-implantation development. Individual data points are overlaid on boxplots for each stage. Differential expression analysis was performed using DESeq2, comparing consecutive developmental stages. Significant log2 fold changes are annotated above comparisons: 4C vs 8C shows significant downregulation (log2FC = −0.84, padj = 0.0301), while 8C vs morula shows significant upregulation (log2FC = 1.7, padj = 0.0235). Non-significant comparisons include 1C vs 2C (log2FC = 1.12, padj = 0.0883), 2C vs 4C (log2FC = 0.174, padj = 0.925), and morula vs blastocyst (log2FC = −0.95, padj = 0.361).

**Table 1:** Datasets used in the study.

| Spermatogenesis |  |  |  |  |
| --- | --- | --- | --- | --- |
| GEO accession | scRNAseq | Before filtering | After filtering | Total |
|  |  | Genes * Cells | Genes * Cells |  |
| GSE109037 [61] | Dataset1 | 33694 * 11104 | 28320 * 10162 | 45067 * 46027 |
|  | Dataset2 | 33694 * 4884 | 27405 * 4628 |  |
|  | Dataset3 | 33694 * 7434 | 27898 * 7029 |  |
|  | Dataset4 | 33694 * 7134 | 29119 * 6665 |  |
| GSE106487 [62] | Dataset5 | 21761 * 3046 | 21761 * 2893 |  |
| GSE153947 [63] | Dataset6 | 56981 * 5437 | 30008 * 5162 |  |
|  | Dataset7 | 56981 * 5438 | 32876 * 5148 |  |
|  | Dataset8 | 56981 * 4672 | 32357 * 4340 |  |
| Testis dataset |  |  |  |  |
| GSE74896 [64] | RNAseq (bulk) |  |  | 12 Samples |
| Human Oocytes |  |  |  |  |
| GSE95477 [65] | RNAseq (bulk) |  |  | 20 samples |
| GSE124718 [27] | CUT&RUN, ATAC-seq |  |  |  |
| Human Pre-Implantation Embryos |  |  |  |  |
| GSE190544 [25] | RNAseq (bulk) |  |  | 73 samples |
| SRP163205 [26] | ATACseq |  |  |  |
| GSE124718 [27] | CUT&RUN, ATAC-seq |  |  |  |
| GSE36552 [55] | scRNAseq |  |  | 124 cells (human pre-implantation embryos and human embryonic stem cells) |
| PGCs (human embryos) |  |  |  |  |
| GSE63818 [51] | scRNAseq | 15 embryos (4 and 19 weeks of gestation) | 233 individual male and female human PGCs, 86 neighbouring somatic cells from 13 of these embryos | 19200 * 328 |

At the 1C (Figure 3a) and 2C (Figure 3b) stages, RNA-seq coverage at the THAP9 locus was minimal, and chromatin accessibility (ATAC-seq) remained low, consistent with a transcriptionally silent state. By the 4C stage (Figure 3c), THAP9 transcripts became detectable, coinciding with strong promoter-associated H3K4me3 enrichment despite only modest chromatin accessibility. This suggests that transcriptional activity at this stage was primarily supported by promoter activation rather than extensive enhancer engagement.

**Figure 3:**
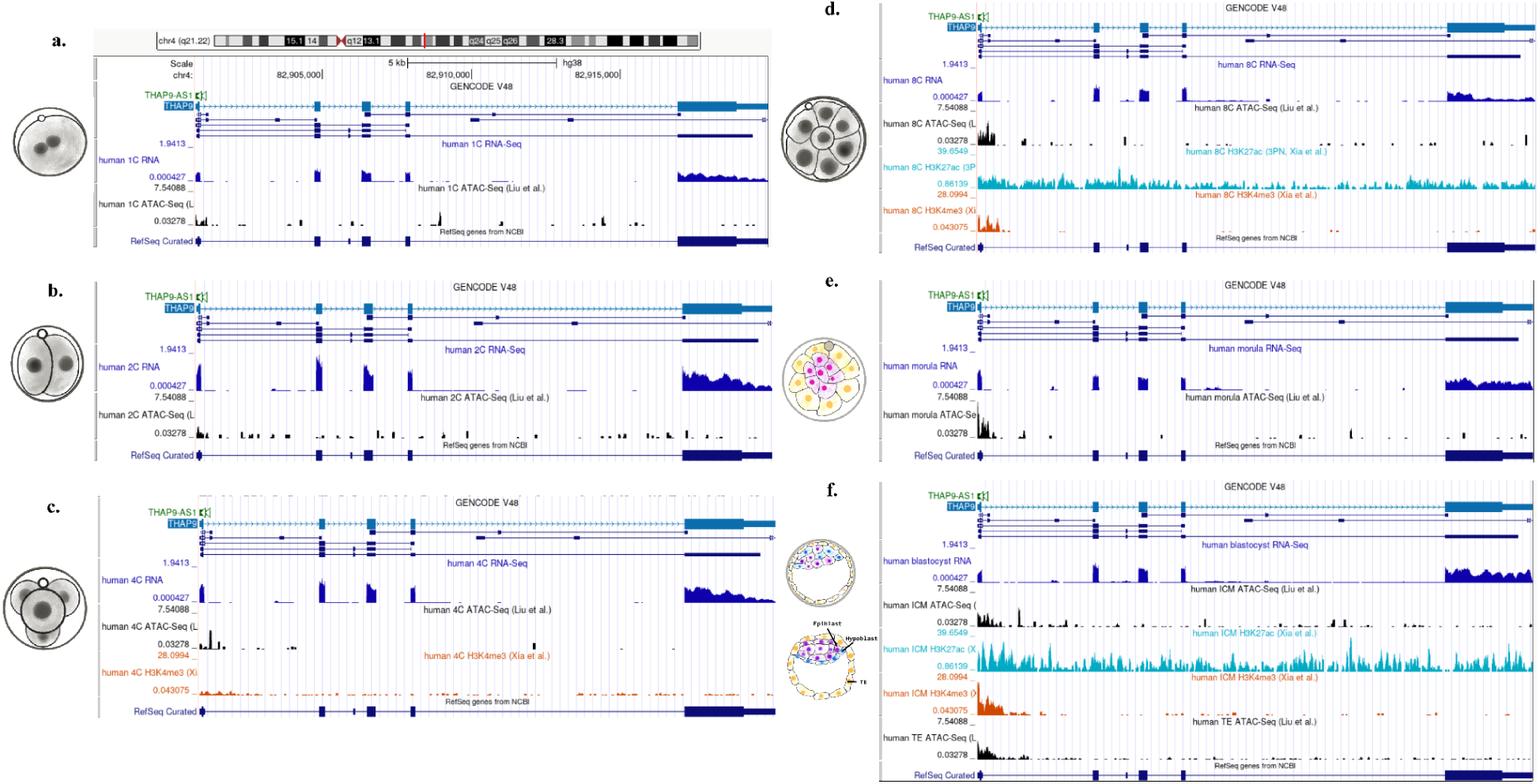
Developmental stage-specific chromatin dynamics at the THAP9 locus during human preimplantation development. UCSC Genome Browser (http://genome.ucsc.edu) views of the THAP9 genomic region (chr4:82,905,000-82,915,000; GRCh38/hg38) showing multi-omic profiles across six developmental stages: **(a)** 1-cell (1C), (**b)** 2-cell (2C), **(c)** 4-cell (4C), **(d)** 8-cell (8C), **(e)** morula, and **(f)** blastocyst (ICM, TE). Each panel displays RNA-seq expression data [25], ATAC-seq chromatin accessibility profiles [26] alongside GENCODE v48 gene annotations. Schematic diagrams on the left illustrate the corresponding embryonic stages.

Between the 4C and 8C stages, THAP9 expression decreased significantly (log2FC = –0.84, padj = 0.0301) (Figure 2b). Paradoxically, epigenomic profiling at 8C (Figure 3d) revealed a gain in chromatin accessibility and acquisition of the enhancer-associated mark H3K27ac, alongside persistent promoter H3K4me3. This indicates that the locus remained epigenetically active and transcriptionally competent, despite reduced RNA abundance. Such a configuration is consistent with a transient transcriptional pause or post-transcriptional regulation [28,29] rather than promoter silencing, keeping THAP9 in a poised state.

At the morula stage, THAP9 underwent robust reactivation (8C to morula: log2FC = 1.70, padj = 0.0235) (Figure 2b), with RNA-seq showing maximal expression and sustained ATAC-seq peaks across both promoter and putative enhancer regions (Figure 3e). From morula to blastocyst, expression decreased modestly (log2FC = –0.95, padj = 0.361) (Figure 2b), though this change was not statistically significant. Chromatin accessibility and histone modification patterns (Figure 3f) at this stage displayed lineage-specific differences: the inner cell mass (ICM) retained strong H3K27ac enrichment and moderate accessibility, whereas the trophectoderm (TE) exhibited more restricted chromatin opening and weaker histone modification signals. The data reveal temporal regulation of THAP9 transcriptional activity during early human embryogenesis coincides with dynamic chromatin remodelling and chromatin accessibility, and active histone marks.

To characterise the epigenetic regulation of THAP9 during early mammalian embryogenesis, we analysed the distribution of key histone modifications across different developmental stages from the 2-cell (2C) stage through blastocyst formation. We examined the active chromatin mark H3K4me3, the repressive mark H3K27me3, and the active enhancer/promoter mark H3K27ac at the THAP9 locus. At the 2C stage (Figure 4a), THAP9 exhibited a configured chromatin configuration characterised by the occurrence of repressive H3K27me3 marks across the gene body and regulatory regions. During the 4C stage (Figure 4b), we observed a dynamic shift in the chromatin landscape of THAP9. The H3K4me3 signal (Figure 4b) became more broadly distributed with increased intensity at the promoter region, while H3K27me3 marks (Figure 4b) showed a more restricted pattern compared to the 2C stage (Figure 4a). This transition suggests the beginning of transcriptional priming at the THAP9 locus. At the 8C stage (Figure 4c), THAP9 displayed further chromatin remodelling with enhanced H3K4me3 deposition at the transcription start site and gene body regions. The H3K27me3 signal became more localised, while H3K27ac marks showed increased abundance, particularly at putative enhancer regions, indicating active transcriptional regulation. In blastocyst-stage embryos, THAP9 exhibited distinct chromatin signatures between the inner cell mass (ICM) and trophectoderm (TE) lineages. In ICM cells (Figure 4d), H3K4me3 and H3K27me3 maintained a balanced distribution with moderate H3K27ac enrichment, consistent with pluripotent cell characteristics, indicating that the gene is transcriptionally poised. A transcriptionally poised state also known as bivalent state, is characterized by the presence of both activating (H3K4me3) and repressive (H3K27me3) histone marks, which keeps them silent yet ready for rapid activation upon lineage commitment [32]. Such poised states are important during early embryonic development, as they allow pluripotent cells to flexibly and precisely initiate lineage specific gene expression in response to differentiation [30].

**Figure 4:**
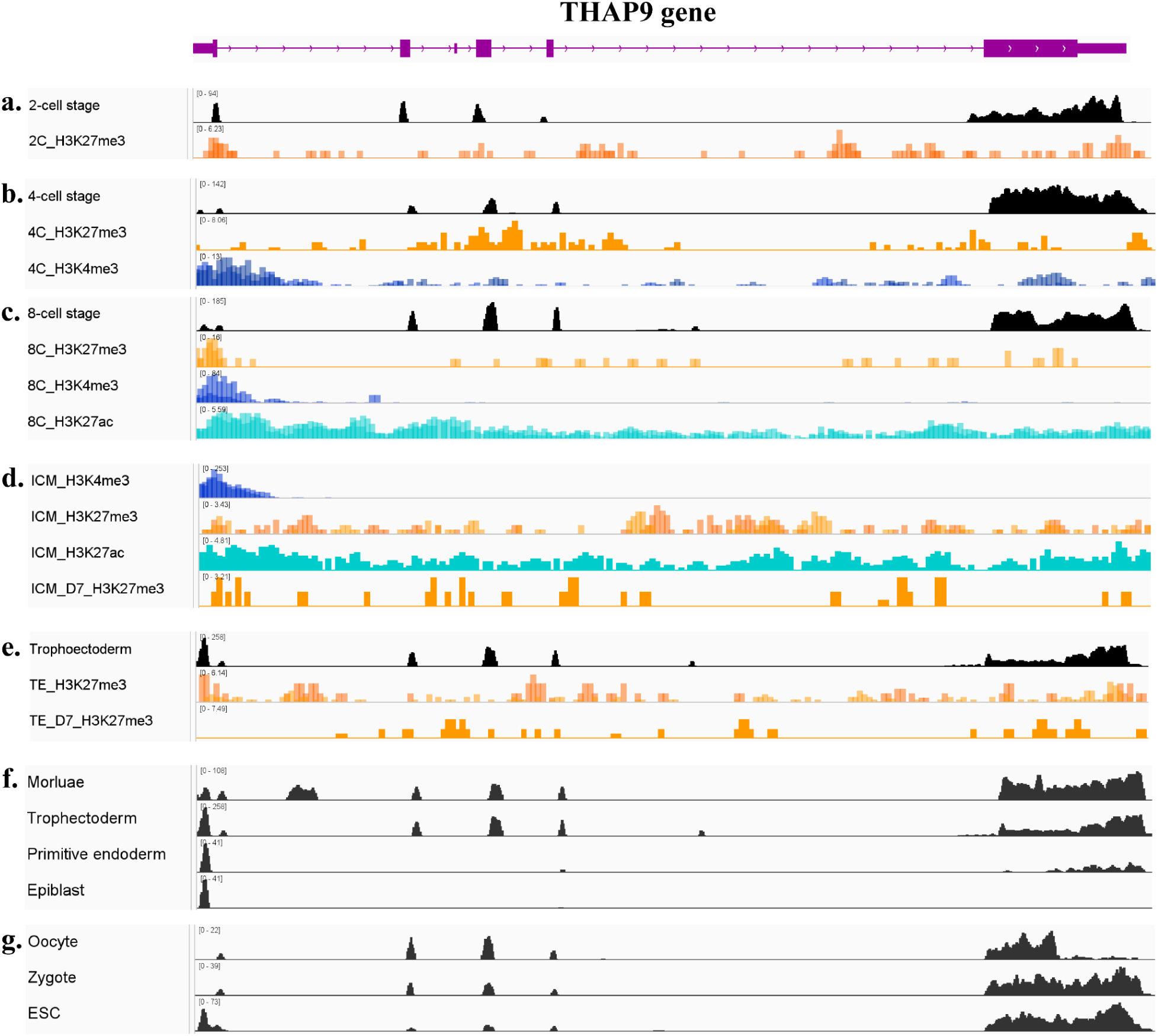
Dynamic chromatin modification and transcriptional activation at the THAP9 locus during early human embryonic development: IGV tracks showing histone modification and expression profiles across the THAP9 gene locus from the oocyte to embryonic stem cell (ESC) stages. (a–e) CUT&RUN tracks illustrate enrichment of H3K27me3 (orange, Polycomb-mediated repression), H3K4me3 (blue, active promoter mark), and H3K27ac (cyan, enhancer/active promoter mark) at successive developmental stages, including the 2-cell (a), 4-cell (b), 8-cell (c), inner cell mass (ICM; d), and trophectoderm (TE; e) stages. (f–g) RNA-seq tracks (black) depict corresponding THAP9 transcript levels at single cell level across the morula, trophectoderm, primitive endoderm, epiblast, oocyte, zygote, and ESC stages. The data highlight dynamic transitions in repressive and active histone marks accompanying transcriptional activation of THAP9 during preimplantation development.

By day 7, H3K27me3 levels decreased, suggesting resolution of bivalency and a shift toward transcriptional activation. In contrast, the trophectoderm (TE) (Figure 4e) THAP9 was predominantly enriched for H3K27me3, consistent with Polycomb-mediated repression, whereby Polycomb complexes silence target genes via chromatin compaction and repressive histone modifications[33]. By day 7, H3K27me3 distribution became more focal and reduced in intensity, indicative of a partial release of repression but without strong evidence of activation. Together, these findings demonstrate that THAP9 is maintained in a poised state in pluripotent ICM cells but is repressed in TE, highlighting its potential role in lineage-specific regulation during blastocyst development.

These findings demonstrate that THAP9 undergoes dynamic transcriptional regulation during preimplantation development, with coordinated chromatin remodelling supporting stage-specific expression patterns that may contribute to zygotic genome activation and early lineage specification (Figure 4f, g). THAP9 undergoes progressive chromatin remodelling during early embryogenesis, transitioning from a bivalent state in early cleavage stages to lineage-specific chromatin configurations in the blastocyst, reflecting its potential role in cell fate specification during mammalian development.

### THAP9 is selectively enriched in spermatids and elongated spermatocytes during human spermatogenesis

Having established the role of the THAP9 gene in early germ cell development, we next investigated its expression during adult male gametogenesis. Analysis of adult tissue-level transcriptomic profiles from the GTEx portal confirmed that THAP9 is most strongly expressed in the human testis (Suppl. Fig.1). To resolve the cellular origin of this expression, we analysed bulk RNA-seq data (Table 1), which included Leydig cells, peritubular cells, Sertoli cells, spermatocytes, spermatids, and whole testis. This revealed that THAP9 expression was predominantly restricted to spermatids, with minimal signal in somatic compartments of the testis (Figure 5a).

**Figure 5:**
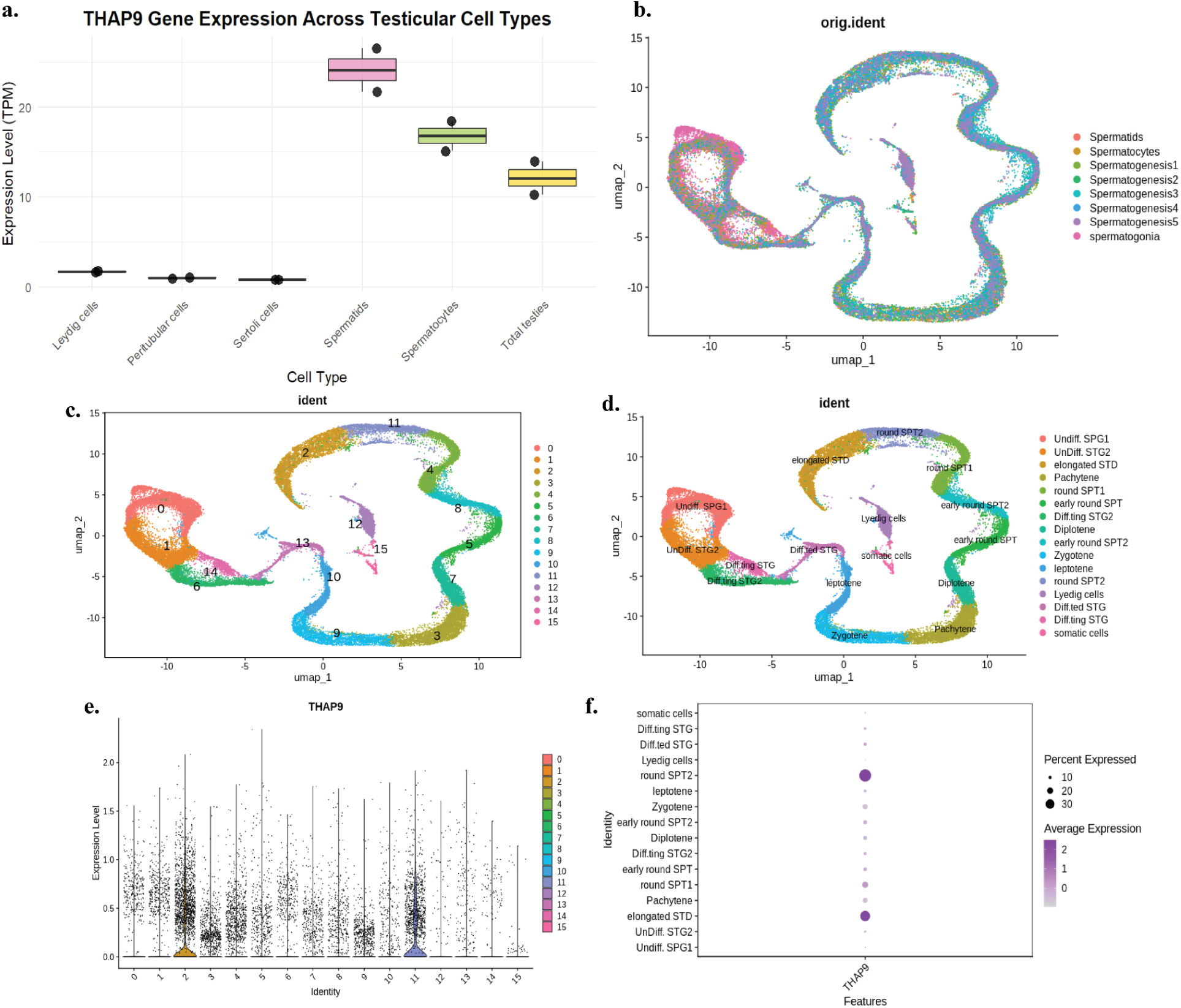
THAP9 gene expression analysis in testicular cells using bulk and single-cell RNA sequencing: **(a)** Bulk RNA-seq analysis showing THAP9 expression levels (TPM) across different testicular cell types. **(b)** UMAP visualisation showing data integration of single-cell RNA-seq datasets (orig.ident), demonstrating successful integration of different experimental datasets **(c)** UMAP plot with numbered cluster identities (0-15) corresponding to different cell populations identified through unsupervised clustering of the single-cell data. **(d)** Annotated UMAP showing detailed cell type classifications including undifferentiated spermatogonia (SPG1, STG2), elongated spermatids (STD), pachytene and round spermatids (SPT1, SPT2), differentiating spermatogonia (STG), diploid cells, zygotene cells, leptotene cells, Leydig cells, and somatic cell populations. **(e)** Violin plot displaying THAP9 expression distribution across the 16 identified cell clusters, showing variable expression levels with some clusters exhibiting higher expression than others. **(f)** Dot plot summarising THAP9 expression across different testicular cell types, where dot size represents the percentage of cells expressing the gene and colour intensity indicates average expression level.

To delineate THAP9 expression at higher resolution, we integrated eight single-cell RNA-seq datasets (Figure 5b) of human spermatogenesis from three independent studies (Table 1), generating an atlas of 46,027 high-quality cells spanning the full germline differentiation continuum (Suppl. Fig. 2a). Clustering and annotation using marker genes (Figure 5c,d) identified spermatogonia (n = 10,162), spermatocytes (n = 4,628), spermatids (n = 7,029), and intermediate germ cell stages (n = 2,893–6,665 each), recapitulating spermatogenic progression (Suppl. Fig. 2b).

Within this framework, THAP9 expression was significantly enriched in round spermatids (cluster 2; log2FC = 1.79, p < 0.001) and elongated spermatocytes (cluster 11; log2FC = 1.67, p < 0.001) (Figure 5e,f). The proportion of THAP9-positive cells in these clusters (32.7% and 39.1%, respectively) exceeded those in other germ cell types (12.3–13.0%). Pseudotime trajectory analysis confirmed that these expression peaks aligned with late stages of spermatogenic progression, validating the developmental annotation of these clusters and demonstrating that THAP9 functions during terminal stages of male germ cell differentiation (Suppl. Fig. 2c,d). THAP9 showed significant upregulation in spermatogenic cell clusters, with the highest enrichment observed in post-meiotic germ cells, particularly round and elongated spermatids.

To explore whether this expression pattern is evolutionary conserved across primates, we examined single-nucleus RNAseq (snRNAseq) data as reported by Mural et al. *[34]* from five primate species, including *Homo sapiens, Pan troglodytes, Gorilla gorilla, Nomascus leucogenys,* and *Macaca mulatta*. UMAP embeddings of all testicular nuclei revealed distinct clustering by annotation of cell type. Visualisation of normalised THAP9 expression across these embeddings showed cell type-specific enrichment (scale 0-3), with binned expression histograms indicating that THAP9 transcripts were predominantly detected in post-meiotic germ cells (Figure 6). This consistent enrichment across all five primate species (Figure 6) supports a conserved post-meiotic (spermiogenic) expression pattern, suggesting that THAP9 could be involved in the functions that are predominantly involved during the spermatid maturation and spermiogenesis, rather than any direct involvement in mitotic spermatogonia or meiotic spermatocytes.

**Figure 6:**
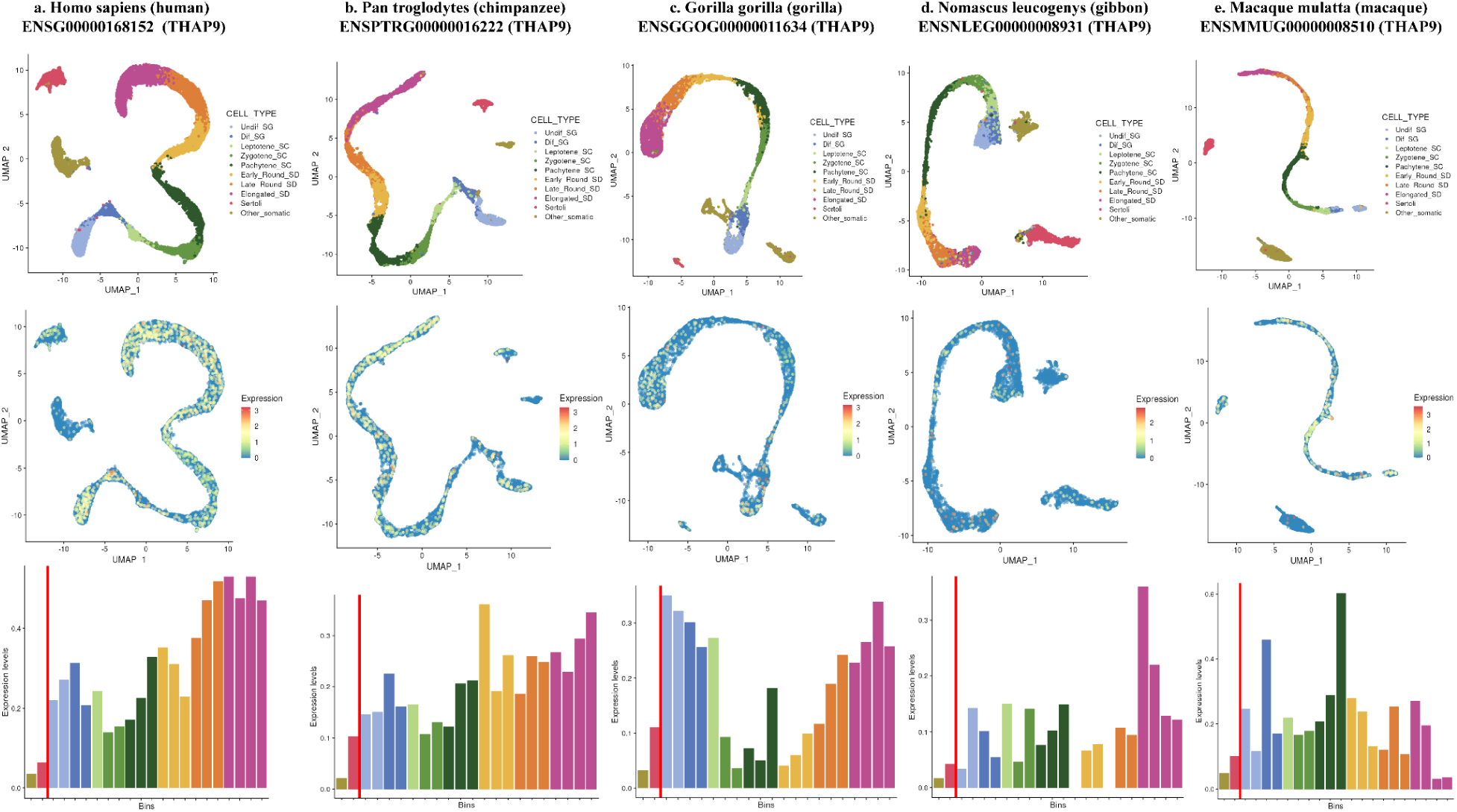
THAP9 expression across spermatogenic cell types in primates (snRNA-seq): Panels show single-nucleus RNA-seq (snRNA-seq) data [34] from five primate species (columns a–e: Homo sapiens, Pan troglodytes, Gorilla gorilla, Nomascus leucogenys, Macaca mulatta), plotted from the dataset published in Murat et al., Nature (2023). Top row: UMAP embedding of all testis nuclei coloured by annotated cell type (legend at right). Middle row: the same UMAP coordinates with dots coloured by normalised THAP9 expression (scale bar: 0–3). Bottom row: binned expression histograms showing the distribution of THAP9 expression across annotated cell-type bins (bars coloured to match cell-type palette); the red vertical line indicates the bin containing the median expression across all nuclei. (Figure created using web tool: https://apps.kaessmannlab.org/SpermEvol/)

### THAP9 associates with a spermatid-specific co-expression program

To explore the transcriptional context of THAP9 expression during spermatogenesis, we constructed co-expression networks using hdWGCNA (Figure 7a). THAP9 was assigned to the spermatid-enriched M7 (yellow) module (eigengene correlation r = 0.276), which represents a transcriptional program strongly associated with spermatid maturation. The hub gene network of this module encompassed genes playing important roles in spermatogenesis and male fertility through diverse mechanisms (Figure 7b). PRSS37 is required for sperm function via ADAM3 maturation, and its deficiency causes male infertility in mice and is linked to unexplained infertility in men [35,36]. CST8, aka CRES, regulates capacitation [37] and the acrosome reaction, with knockout mice showing impaired in vitro fertilisation [38]. IFT172 and TEKT5 are essential for flagellar assembly and motility [39,40]. MAPK3 (ERK1) supports spermatogonial proliferation and sperm functions, with pathway dysregulation linked to subfertility [41]. CAP1 loss in hamsters results in mid-piece loss in spermatozoa [42]. PRSS58 and REXO1 are enriched in spermatids or testis transcriptomics but lack direct functional or disease evidence [43,44]. These highlight the role of the THAP9 gene in late germ cell specialisation (Figure 6b).

**Figure 7:**
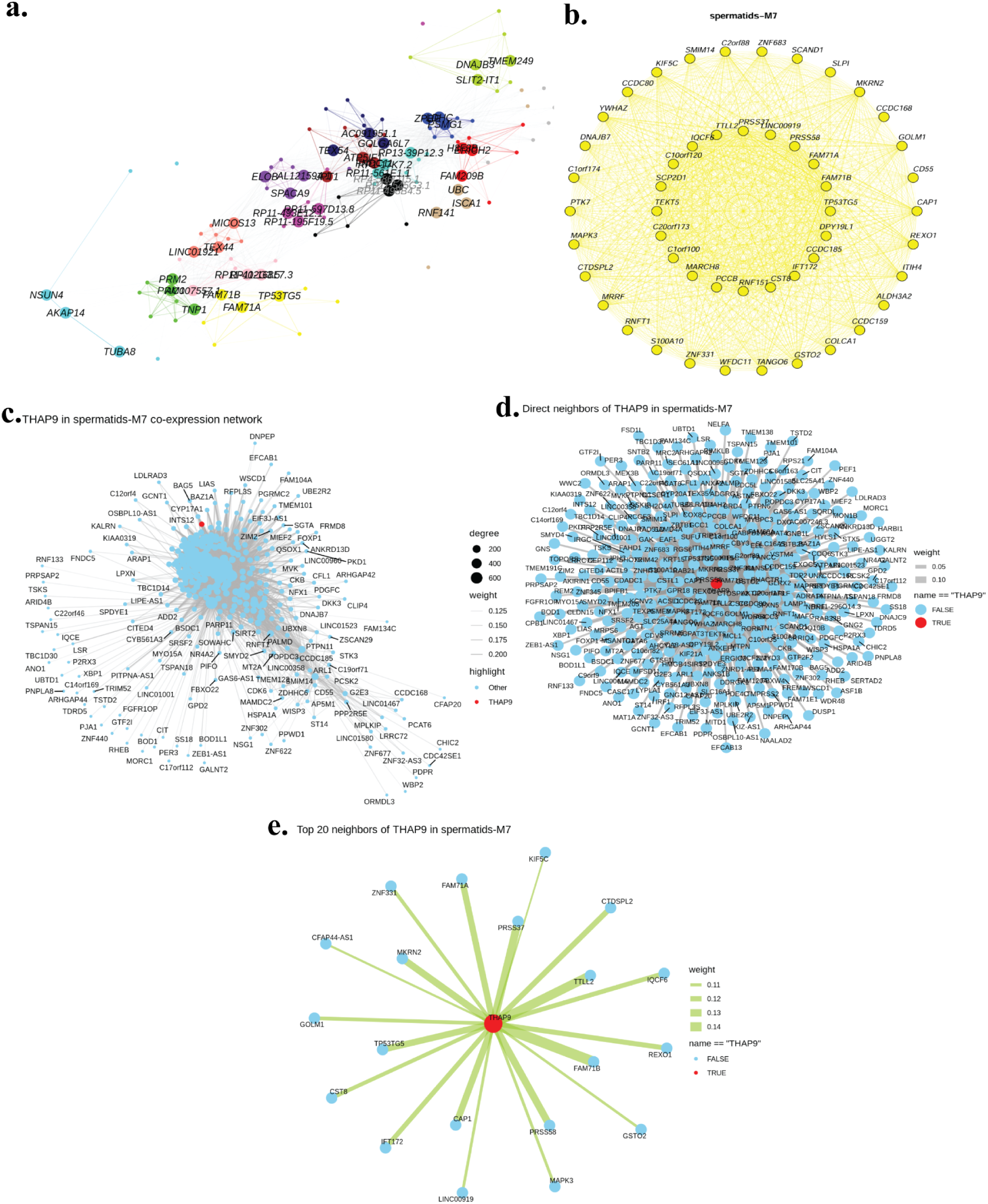
Co-expression network analysis of THAP9 in spermatids-M7 using hdWGCNA. **(a)** Hub network visualisation showing the overall network structure with major hub genes and their connections. **(b)** Complete co-expression network highlighting the yellow module where THAP9 is located. Yellow nodes represent genes within the spermatids-M7 module. **(c)** The THAP9 gene (highlighted in red) is positioned within the yellow module co-expression network, showing its integration within the module structure. **(d)** Direct neighbour network of THAP9 showing all genes with direct co-expression connections to THAP9. Node size reflects degree centrality, edge weight represents co-expression strength, and the network layout emphasises THAP9’s central position (red node) among its immediate neighbours. **(e)** Top 20 direct neighbours of THAP9 ranked by co-expression weight. THAP9 (red node) is shown at the centre with its strongest co-expressed genes connected by edges weighted according to co-expression strength (edge thickness and colour intensity indicate relationship strength). Analysis performed using the hdWGCNA package to identify co-expression modules and gene relationships in spermatids-M7.

Within this network, THAP9 was embedded among spermatid-associated hub genes (Figure 7c). To refine its functional associations, we extracted its direct neighbours, identifying genes that were direct co-expression partners of the THAP9 gene (Figure 7d). From this set, the top 20 most strongly connected genes (Figure 7e) included testis-enriched factors involved in spermatogenesis (PRSS37, PRSS58, IQCF6, CST8), ciliary/flagellar function (IFT172, KIF5C, CFAP44-AS1), and signalling regulation (MAPK3, CAP1). Functional enrichment analysis of these neighbours revealed overrepresentation of pathways linked to spermatid differentiation, cilium organisation, and sperm motility.

Together, these findings demonstrate that THAP9 participates in a spermatid-specific co-expression program, aligning with transcriptional modules that regulate terminal germ cell differentiation and structural maturation, consistent with its expression pattern during embryonic germ cell development.

### THAP9 shows stage-specific regulation in human oocytes and associates with nuclear transport and cell cycle networks

To determine whether THAP9 also contributes to female gametogenesis, we analysed transcriptome data of immature (germinal vesicle, GV), which are arrested at prophase I of meiosis, mature (metaphase II, MII) human oocytes (Figure 8a), which are arrested at metaphase II of meiosis, from young (<30 years) and advanced maternal age (≥40 years) patients (Table 1). Principal component analysis (Figure 8b) clearly separated GV and MII oocytes, and differential expression analysis identified genes with significant changes between stages (Figure 8c), including THAP9 gene expression, which is significantly higher in germinal vesicle oocytes compared to metaphase II oocytes, with a Log2 fold change of 1.58 and an extremely significant adjusted p-value of 1.03e-25 (Figure 8d). The data show THAP9 expression decreases from germinal vesicle stage oocytes to metaphase II oocytes.

**Figure 8:**
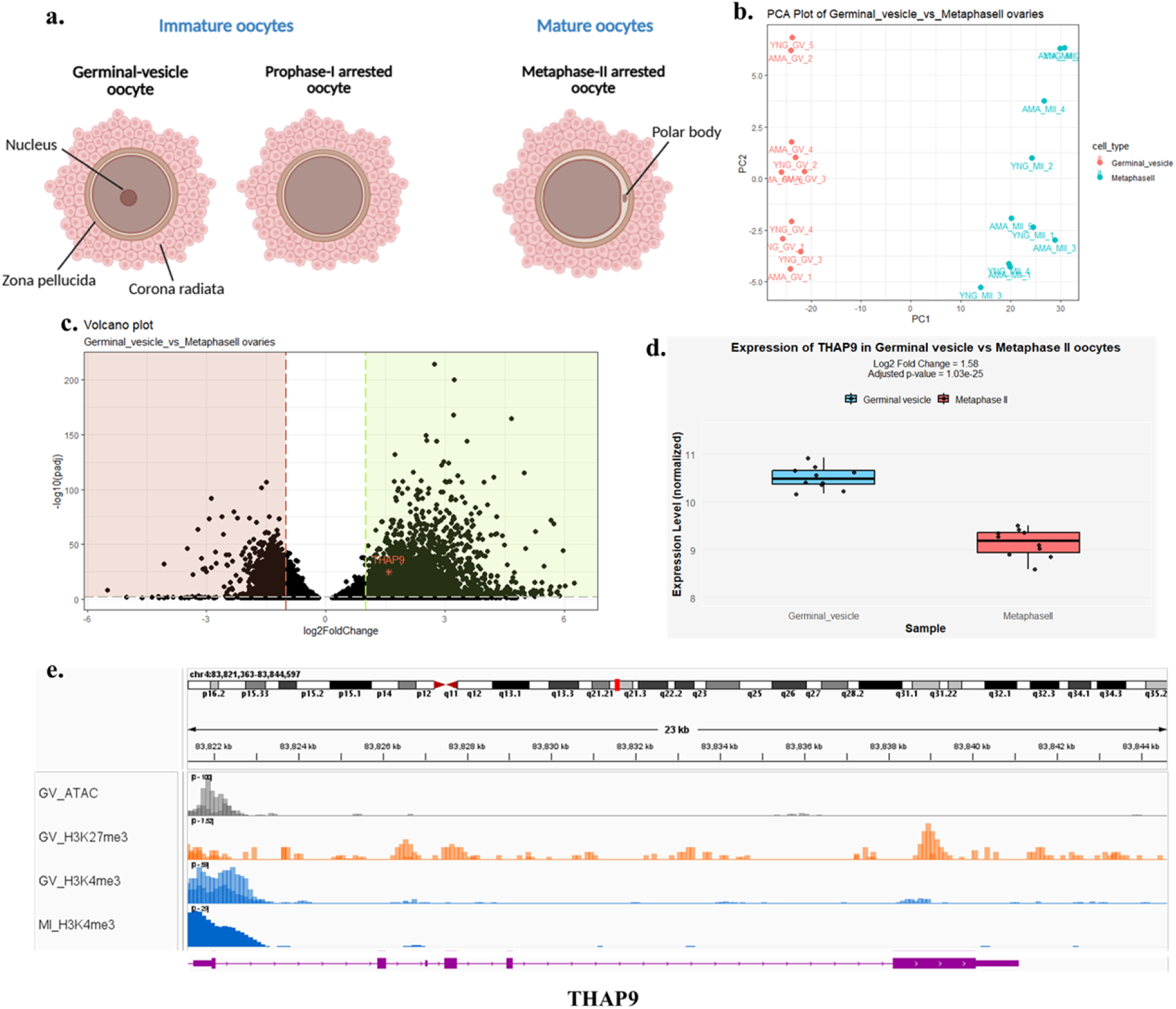
THAP9 expression is stage-specifically regulated during human oocyte maturation through coordinated transcriptional and epigenetic mechanisms. **(a)** Stages of human oocyte maturation: Representative diagram showing the progression from immature to mature oocytes. Immature stages include the germinal-vesicle (GV) oocyte and the meiosis-I arrested oocyte (MI); the mature stage corresponds to the Metaphase-II (MII) arrested oocyte. (**b)** Principal component analysis (PCA) of germinal vesicle (GV, red) and metaphase II (MII, cyan) oocytes from young and advanced maternal age patients, showing clear transcriptional separation between maturation stages. **(c)** Volcano plot of differential gene expression between GV and MII oocytes. THAP9 is highlighted in red. Pink and green shaded regions indicate genes which are downregulated and upregulated. **(d)** Box plot comparing THAP9 expression levels between GV and MII oocytes, confirming significantly higher expression in immature (GV) compared to mature (MII) oocytes. **(e)** Epigenomic landscape at the THAP9 locus showing chromatin accessibility (ATAC-seq, grey tracks) and histone modifications (H3K27me3, orange tracks and, H3K4me3, blue tracks) in GV and MI oocytes. GV oocytes show active chromatin marks (H3K4me3, blue tracks) and open chromatin accessibility at the THAP9 promoter, while MI oocytes exhibit active promoter marks (H3K4me3). The genomic coordinates and gene structure are shown at the top.

We examined chromatin accessibility and histone modification patterns at the THAP9 locus (Figure 8e). In GV oocytes, the THAP9 promoter region showed strong enrichment for the active transcription mark H3K4me3 and open chromatin accessibility (ATAC-seq), consistent with active transcription. However, the repressor-associated mark H3K27me3 was widespread, suggesting the signal of repression; H3K4me3 is a “counter-signal” at the promoter, but it hasn’t overridden the repression for full activation. MI oocytes exhibited higher H3K4me3 signals at the promoter (5’ end) region, suggesting an active promoter in meiosis-I stage oocytes, which is meiosis-I arrested oocyte.

To understand the functional relationship of this gene, co-expression network analysis was performed using WGCNA with a soft-thresholding power of 10 (Figure 9a). THAP9 was assigned to the turquoise module, which was significantly correlated with GV-stage oocytes (Figure 9b). Notably, THAP9 exhibited a high module membership (MM = 0.98), indicating that it is a strongly representative gene of this expression program (Figure 9c,d). The turquoise module network (weight > 0.4) highlighted THAP9 as a central node connected to multiple genes involved in RNA processing, nuclear transport, and chromatin organisation (Figure 9e). Extraction of the top 10 most strongly co-expressed partners identified genes such as KPNA2, CCT3, CCT5, KIF22, and HMGN2, which have established roles in nuclear import, chromatin remodelling, and spindle assembly (Figure 9f) [45–49] These findings reveal that THAP9 is dynamically regulated during oocyte maturation through coordinated transcriptional and epigenetic mechanisms, showing high expression in GV oocytes but repression at the MII stage, in contrast to its upregulation during spermatid differentiation. This sex-specific difference in temporal regulation suggests distinct functional requirements for THAP9 in male versus female gametogenesis. Moreover, its strong co-expression with nuclear transport and cell cycle regulators suggests a potential role in transcriptional control and nuclear remodelling events unique to oocyte maturation, complementing its structural roles observed during spermatogenesis.

**Figure 9:**
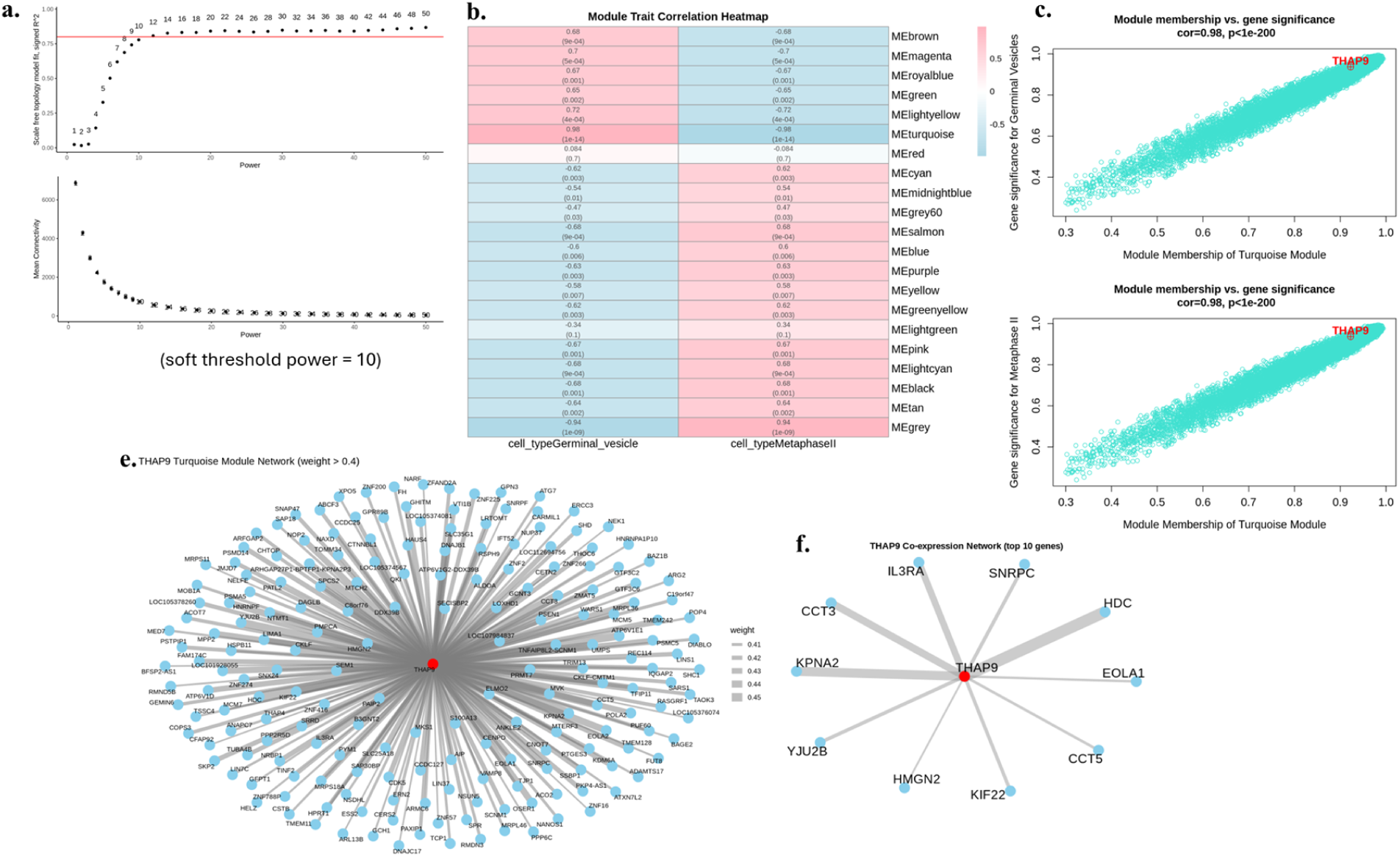
Weighted Gene Co-expression Network Analysis (WGCNA) based Identification of THAP9 and Its Co-expressed Gene Network. **(a)** Analysis of network topology to determine soft-thresholding power. A power of 10 was chosen as the optimal soft threshold to achieve scale-free topology. **(b)** Module–trait correlation heatmap showing associations between gene co-expression modules and oocyte developmental stages (germinal vesicle and metaphase II). The turquoise module shows the strongest positive correlation with both stages. **(c–d)** Scatterplots of module membership versus gene significance for the turquoise module, highlighting THAP9 as a highly significant hub gene strongly correlated with germinal vesicle (c) and metaphase II (d) stages. **(e)** Gene network of the turquoise module (edge weight > 0.4), showing THAP9 (red) as a central hub gene connected to multiple co-expressed partners. **(f)** Subnetwork of THAP9 with top co-expressed genes

## Discussion

In this study, we provide the first integrated view of THAP9 transcriptional landscape and potential functions across the germline and early embryonic stages. Our analysis provides a unifying theme suggesting that THAP9 is engaged during transcriptional reprogramming and terminal differentiation, processes that require extensive chromatin remodelling and establishment of new cell identities. We systematically investigated the developmental and germline regulation of the THAP9 gene, combining bulk and single-cell transcriptomic datasets with epigenomic profiling. By integrating data spanning human embryogenesis, gametogenesis, and preimplantation development, we provide the first comprehensive characterisation of THAP9 expression dynamics and chromatin regulation.

We show that THAP9 is strongly enriched in PGCs during early development, progressively silenced in somatic lineages, dynamically regulated during oogenesis and spermatogenesis, and subjected to coordinated chromatin remodelling during preimplantation development. These findings collectively identify THAP9 as a germline-enriched and developmentally regulated gene with potential importance in early human reproduction. During embryogenesis, THAP9 was consistently enriched in PGCs compared with somatic cells, with maximal expression between developmental weeks 7 and 11. This window corresponds to germline migration and gonadal colonisation, an important period for establishing the germline program [50,51]. In contrast, somatic lineages showed progressive downregulation, suggesting that THAP9 regulation is germline-specific and temporally restricted. This selective enrichment parallels other germline-restricted regulators such as SOX17 and TFAP2C, which safeguard PGC identity [52,53].

In the adult testis, THAP9 expression was most prominent in spermatids, with single-cell resolution analysis showing enrichment in round spermatids and elongated spermatocytes. Co-expression network analysis further embedded THAP9 in a transcriptional module enriched for genes regulating acrosome biogenesis, flagellar assembly, and cytoskeletal remodelling. This positions THAP9 alongside established regulators of terminal germ cell differentiation and sperm maturation. This supports a role for THAP9 in transcriptional programs governing terminal spermatogenic differentiation and sperm maturation. In contrast, in the ovary, THAP9 expression was highest in germinal vesicle (GV) oocytes and downregulated in metaphase II (MII) oocytes. Network analysis positioned THAP9 among genes associated with nuclear transport, chromatin organisation, and spindle assembly. These findings suggest that THAP9 contributes to nuclear and transcriptional remodelling in early oocyte maturation, in contrast to its later functional timing in spermatogenesis. Interestingly, the second-highest expression (Suppl Fig. 1) is observed in the thyroid, an endocrine organ that plays a key regulatory role in reproductive function, further supporting a link between THAP9 activity and reproductive physiology. Together, these results highlight distinct temporal requirements for THAP9 in male versus female gametogenesis.

Our analysis of pre-implantation embryos further highlights the dynamic regulation of THAP9 during early human development. THAP9 expression was minimal at the zygote 4 cell stages with a marked increase from the 8 cell stages to the morula, coinciding with major zygotic genome activation (ZGA) [54,55]. The transition from bivalent to more permissive chromatin states between the 4-cell and 8-cell stages, marked by enhanced H3K4me3 deposition and the acquisition of H3K27ac at putative enhancer regions, indicates progressive chromatin opening and transcriptional priming. Notably, the apparent paradox observed at the 8-cell stage, where THAP9 expression decreases despite increased chromatin accessibility and enhancer activation, suggests sophisticated post-transcriptional regulatory mechanisms. This configuration may represent a transcriptional pause that maintains the locus in a poised state, ready for the robust reactivation observed at the morula stage. Interestingly, a transient downregulation occurred at the 8-cell stage despite increased chromatin accessibility and enhancer-associated H3K27ac enrichment. This indicates that THAP9 was maintained in an epigenetically active but transcriptionally retrained state, consistent with transcriptional pausing or post-transcriptional regulation described for other ZGA genes [56]. By the morula stage, THAP9 was robustly reactivated, with strong promoter and enhancer accessibility, and in the blastocyte, regulation became lineage specific; poised in the inner cell mass (ICM) but repressed in the trophoectoderm (TE). Such divergent regulation mirrors pluripotency-associated regulators that remain accessible in ICM while excluded from TE [57–59].

Epigenomic proliferation revealed that THAP9 underwent progressive chromatin remodelling. At early cleavage stages, THAP9 was bivalently marked by H3K4me3 and H3K27me3, a hallmark of genes poised for activation [30,60]. This bivalency was resolved at the morula stage with the acquisition of H3K27ac and robust transcription, while in the blastocyst, ICM retained a poised state, whereas TE exhibits polycomb-mediated repression. The activation of THAP9 at the blastocyst stage, with sustained expression across all three lineages and in human embryonic stem cells (Figure 4f,g), suggests a role in lineage specification and the maintenance of pluripotency. These findings point to THAP9 regulation being integrated into broader chromatin characteristics of lineage specification.

By combining single-cell RNA-seq, bulk tissue transcriptomics, co-expression network analyses, and epigenomic profiling, our work provides a multi-layered map of THAP9 regulation. This integrative bioinformatics approach has been widely applied to characterise developmental regulators, and here it establishes THAP9 as a candidate gene with coordinated transcriptional and epigenomic dynamics in human reproduction. Together, these findings establish THAP9 as a dynamically regulated gene with germline enrichment, sex-specific roles in gametogenesis, and lineage-specific control during early embryogenesis. While the results provide correlative evidence from transcriptomic and epigenomic datasets, future functional studies will be required to define the precise molecular role of THAP9 in human reproduction and development.

## Limitations and Future Directions

This study establishes THAP9 as a dynamically regulated, germline-enriched gene with distinct expression and chromatin states during human pre-implantation development, zygotic genome activation (ZGA), and gametogenesis. The integration of transcriptomic and epigenomic datasets highlights multilayered regulation involving transcriptional activity, chromatin accessibility, and histone modifications, highlighting its potential importance in pluripotency and lineage specification.

Several limitations must be noted. First, the analyses rely exclusively on public transcriptomic and epigenomic datasets, which provide correlative rather than causal evidence of THAP9 function. Future studies employing functional perturbation approaches, such as CRISPR-mediated knockout, knockdown, or overexpression in human embryonic stem cells, induced pluripotent stem cell-derived germ cell models, or organoid systems, will be essential to determine the mechanistic role of THAP9. Second, independent validation across additional datasets and experimental platforms is required to confirm the robustness of these findings. Third, the extent to which regulatory patterns of the THAP9 gene are conserved across species remains unresolved. Comparative studies using non-human primate models, as well as cross-species genomic and epigenomic analyses, will be necessary to determine whether THAP9 represents a human-specific innovation in germline regulation.

Despite these limitations, our study provides a comprehensive framework for investigating how domesticated DNA transposon-derived genes contribute to human development. By highlighting THAP9 as a candidate regulator of pluripotency and germline specification, this work sets the stage for mechanistic studies that could link TE-derived regulators to fertility, developmental disorders, and species-specific aspects of reproduction.

## Methodology

### Bulk RNA-seq data acquisition and processing

Bulk RNAseq data from datasets GSE74896 [64], GSE95477 [65], and GSE190544 [25] were obtained from Gene Expression Omnibus (GEO) repository at National Center for Biotechnology Information (NCBI) [66]. Raw count data and corresponding sample metadata, including stages and other biological variables, were imported into R(v4.4.1) for analysis. Normalised expression values were obtained using the variance-stabilising transformation (VST) implemented in DESeq2(v1.46.0) [67]. Quality control involved principal component analysis (PCA) to identify potential outlier samples and hierarchical clustering based on Euclidean distance with average linkage. Dendrograms were colour-coded according to stages, and sample outliers were inspected and verified manually to ensure data integrity before network analysis. Differential expression analysis was done using DESeq2(v1.46.0) library[67]; the p-values attained by the Wald test were corrected for multiple testing using the Benjamini and Hochberg method [68] by default to obtain adjusted p-values; these corrected p-values were used for analysis and interpretation of results.

### Single-cell RNAseq data processing and integration

Single-cell RNA sequencing (scRNA-seq) data from human testicular tissue were analysed using Seurat(v5.3.0) [69] implemented in R(v4.4.1). Raw count matrices were imported (the datasets reported in Table 1) and preprocessed using Seurat’s standard workflow. Each dataset was log-normalised, and the most variable genes were identified with FindVariableFeatures(). Integration across samples was performed using the anchor-based strategy implemented in Seurat through FindIntegrationAnchors() and IntegrateData(). After integration, the data were scaled, ScaleData(), and dimensionality reduction was conducted using principal component analysis (PCA), RunPCA(). Clustering was performed with the Louvain algorithm (FindClusters()), with the number of principal components set to 30. Dimensionality reduction and visualisation were achieved using Uniform Manifold Approximation and Projection (UMAP), RunUMAP(), with the clustering resolution set to 0.3. Cluster-specific marker genes were identified using FindAllMarkers() and compared with known germ cell markers from the published references to assign biological identities. The annotated Seurat object was visualised using DimPlot() with cell type and sample grouping.

### Trajectory and Pseudotime inference analysis

Developmental trajectory reconstruction and pseudotime inference were performed using Monocle3(v1.4.26) [70] via SeuratWrappers(v0.4.0). The integrated Seurat object was converted to a Monocle3 cell data set (CDS) and processed under the RNA assay, with data layers merged for analysis. Cluster identities and UMAP embeddings from Seurat were transferred to the Monocle3 object to preserve established cell-type relationships. Cells were reclustered, and a trajectory graph was constructed using Monocle3’s learning algorithm without partitioning. Pseudotime ordering was then performed to infer developmental progression, and pseudotime values were added to the Seurat metadata for integrated visualisation. Pseudotime trajectories were visualised using Monocle3 and Seurat plotting functions, and gene expression dynamics along pseudotime were assessed to reveal transcriptional changes across spermatogenic differentiation.

### scRNAseq analysis of PGC data

The dataset from GSE63818 [51] from GEO [66], containing Individual gene expression files, was imported for each sample. Files were imported into Python(v2.7.17) using pandas. Sample metadata, including sex, cell type (PGC or soma), embryonic weeks, embryo ID, and cell ID, were extracted from filenames. Expression matrices were merged into a single matrix with genes as rows and cells as columns. The matrix was transposed to (cells x genes) for downstream analysis. Weeks of gestation were converted to numeric values and filtered for downstream modelling. Pairwise comparison between cell types at each week was performed using Mann-Whitney tests [71]. UMAP visualizations were generated using the Scanpy package [72], with cells colored by their Leiden clustering [73] assignments to illustrate cluster identities and relationships among cells.

### high-dimensional Weighted Gene Co-expression Network Analysis (hdWGCNA)

Single-cell RNA-seq data from male germ cells were analysed using Seurat(v5.3.0). The integrated Seurat object was updated, normalised, and scaled. Cell types were annotated based on prior clustering and verified by UMAP visualisation. Metacells were generated using hdWGCNA(v0.4.05) [74] by grouping cells according to cell type, followed by PCA and Harmony batch correction. Gene filtering retained transcripts expressed in at least 5% of cells. Metacell-level expression matrices were extracted for spermatid populations, and co-expression networks were constructed under a signed configuration using a soft-thresholding power of 10. Modules were identified from the topological overlap matrix (TOM) and renamed for consistency. Module eigengenes were computed in both standard and Harmony-corrected forms, and hub genes were defined as the top 10 genes per module with the highest module membership (kME). The THAP9 gene was assigned to module spermatids-M7 with moderate connectivity (kME = 0.24), and the top 20 THAP9-associated genes were extracted for functional analysis. Module activity across single cells was evaluated using UCell-based scoring. All processed data, including eigengenes, TOMs, and module assignments, were saved to ensure reproducibility.

### Weighted Gene Co-expression Network Analysis (WGCNA)

Co-expression networks were constructed using the Weighted Gene Co-expression Network Analysis (WGCNA)(v1.73) [75] R package. Low-count filtering was performed following WGCNA recommendations, excluding genes with fewer than 15 counts in more than 75% of samples. After filtering, the downstream analysis was performed. To achieve an approximate scale-free topology (R² > 0.8), the soft-thresholding power was determined using the pickSoftThreshold() function under a signed network configuration, and a power of 10 was selected. Network construction and module detection were performed using the blockwise module approach with a maximum block size of 9,000 genes, a minimum module size of 30, a signed TOM, and a merge cut height of 0.25. Module eigengenes were calculated for each module, and their correlations with oocyte developmental stages were assessed using Pearson’s correlation and Student’s t-test. Modules significantly associated with oocyte stages were visualised as dendrograms and heatmaps to confirm the robustness of module assignments. Hub genes were identified based on module membership (MM) and gene significance (GS) within each module. Genes in the turquoise module with |MM| ≥ 0.8 and |GS| ≥ 0.8 were defined as hub genes, among which THAP9 was identified as a key member. The network was exported in Cytoscape-compatible format, and subnetworks centred on hub genes were filtered for edges with weights ≥ 0.4. Visualisation was performed using the igraph(v2.1.4) and ggraph(v2.2.1) R packages, with THAP9 highlighted to illustrate its central role within the turquoise module.

### ATAC-seq and CUT&RUN data

Public ATAC-seq data were obtained from the Sequence Read Archive (SRA), and CUT&RUN data from NCBI’s GEO [66] (Table 1). Raw reads were trimmed and quality-checked using Fastp(v1.0.1) [76], then aligned to the corresponding human reference genome with Bowtie2 (v2.5.4) [77]. Sambamba(v1.0.1) [78] was used to remove low-quality, duplicate, unmapped, and non-canonical chromosomal reads. For ATAC-seq, peaks were identified with Genrich(v0.6.1) (https://github.com/jsh58/Genrich), and for CUT&RUN, MACS3(v3.0.3) [79]. Consensus and differential peaks across developmental stages were defined using DiffBind(v3.18.0). Normalisation factors were estimated using EdgeR(v4.4.2), and scaling factors were applied in deepTools(v3.5.6) [80] bamCoverage(v3.5.6) to generate normalised BigWig tracks. Mean coverage was calculated using wiggletools(v1.2) [81], and BigWig files were visualised in Integrative Genomics Viewer (IGV)(v2.18.2) [82].

### Visualisation

Publicly available datasets were obtained from the GEO [66]. Single-cell transcriptomic profiles of human embryos were downloaded from GSE36552 [55], and histone marks H3K4me3, H3K27me3 and H3K27ac from GSE124718 [27]. Single-cell RNA-seq captures cell-to-cell variability in gene expression, whereas histone marks represent averaged chromatin modification patterns across multiple cells. To visualise and qualitatively compare these data, tracks were loaded into the IGV(v2.18.2)[82] using the hg19 reference genome. This approach enabled the examination of transcriptional activity relative to repressive histone marks in early human embryonic development.

## Data and code availability

No new datasets or custom code were generated in this study.

## Supplementary Figures

**Supplementary Figure 1:**
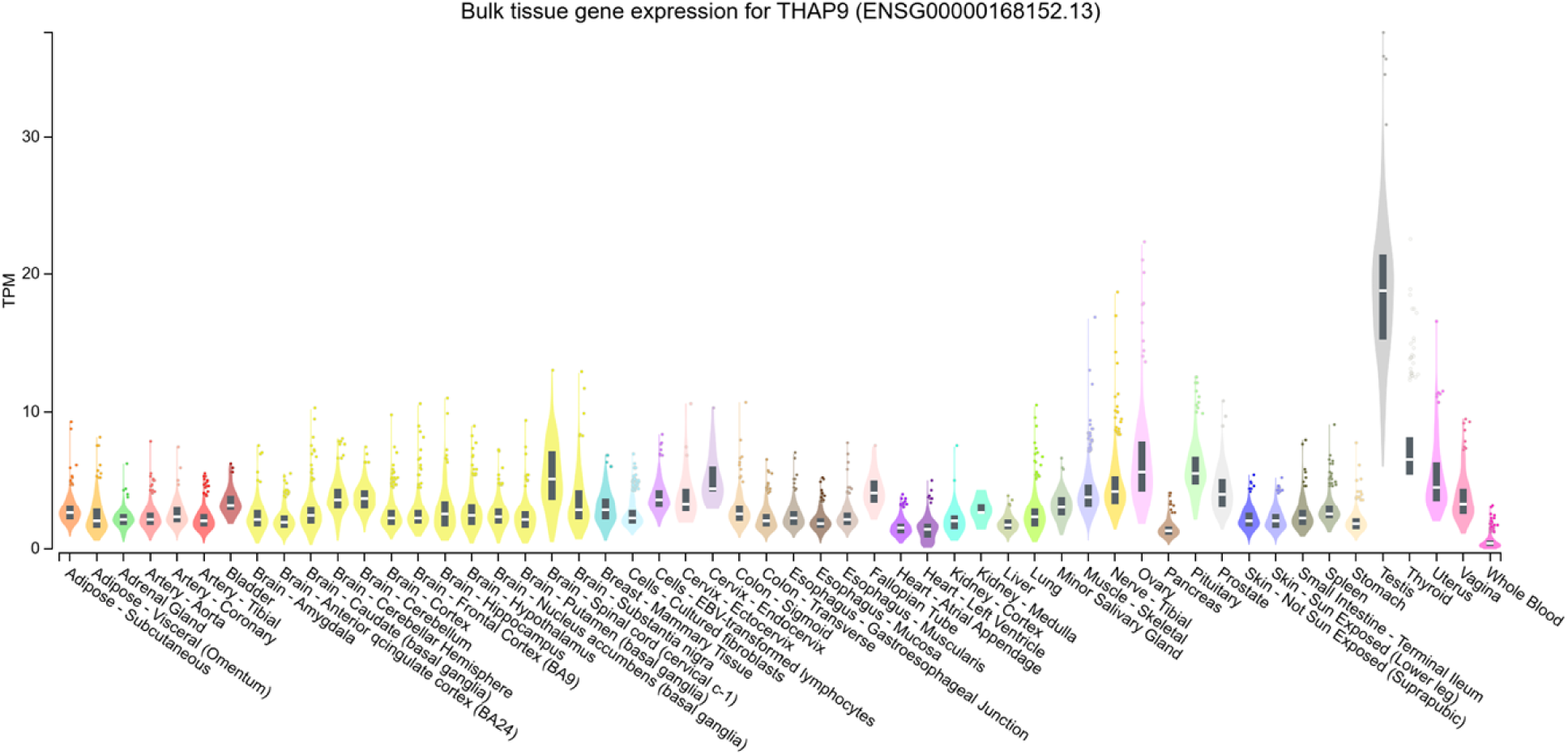
Tissue-specific expression profile of THAP9 gene across human tissues: Violin plots displaying bulk tissue gene expression data for THAP9 (ENSG00000168152.13) showing transcripts per million (TPM) values across diverse human tissues from the GTEx (Genotype-Tissue Expression) portal. The width of each violin represents the density of samples at different expression levels, and black bars within violins indicate median expression values. Data obtained from the GTEx portal (https://gtexportal.org/home/gene/THAP9)

**Supplementary Figure 2:**
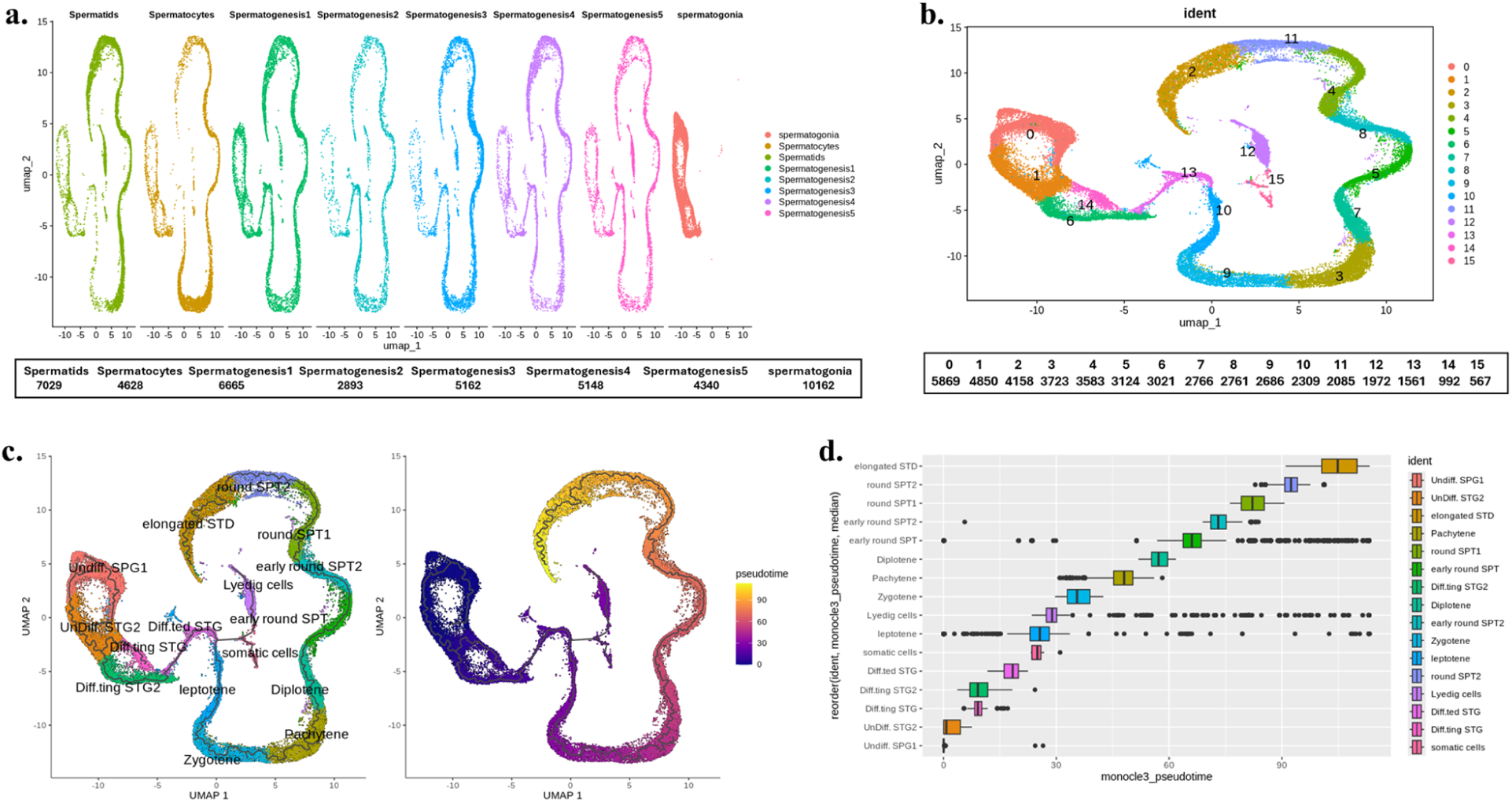
Single-cell RNA sequencing analysis reveals dataset integration, clustering, and developmental trajectory during spermatogenesis. **(a)** UMAP visualisation showing the integration of different spermatogenesis datasets, with each dataset colored separately. Cell numbers for each dataset are indicated below **(b)** UMAP plot displaying 16 distinct cell clusters (0-15) identified through unsupervised clustering of the integrated dataset, with the number of cells in each cluster listed below **(c)** Pseudotime analysis using trajectory inference to reconstruct the developmental progression during spermatogenesis. Left panel shows the inferred trajectory path overlaid on cell populations, while the right panel displays the same trajectory colored by pseudotime values, indicating the temporal ordering of developmental stages **(d)** Hierarchical ordering of cell stages along the developmental trajectory, showing the sequential progression from undifferentiated spermatogonial populations through various differentiated stages including spermatocytes, round spermatids, and elongated spermatids, as determined by pseudotime analysis.

